# AMP-Atlas: Comprehensive Atlas of Antimicrobial Peptides to Combat Multidrug-resistant Bacteria

**DOI:** 10.1101/2025.02.27.640545

**Authors:** Yuki Otani, Daisuke Koga, Yasunari Wakizaka, Hideyuki Shimizu

**Affiliations:** Department of AI Systems Medicine, M&D Data Science Center, Institute of Integrated Research, Institute of Science Tokyo, Tokyo, JAPAN; Graduate School of Medical and Dental Sciences, Institute of Science Tokyo, Tokyo, JAPAN

**Keywords:** Artificial intelligence, Antimicrobial peptides, Drug discovery, Drug-resistant bacteria

## Abstract

The escalating threat of infections caused by drug-resistant bacteria poses a significant global health challenge, with projections estimating 10 million annual deaths by 2050. While the development of conventional antibiotics has stagnated since the late 1990s, antimicrobial peptides (AMPs), short amino acid sequences exhibiting potent antimicrobial activity, have emerged as a promising alternative, demonstrating efficacy even against drug-resistant bacteria. However, despite the identification of numerous AMPs, their translation into clinically approved therapeutics remains limited, highlighting the critical need for accelerated discovery methods that transcend traditional experimental screening. Here, we introduce AMP-Atlas, an AI system inspired by cutting-edge natural language processing, designed to accurately predict antimicrobial activity from peptide sequences alone. AMP-Atlas achieves state-of-the-art performance, outperforming existing methods in AMP identification. Furthermore, we leveraged AMP-Atlas to screen human indigenous bacterial flora species, revealing a vast reservoir of previously unexplored AMP candidates. Our findings underscore the transformative potential of AI-powered approaches to revolutionize AMP discovery and development, paving the way for innovative therapeutic strategies to combat the looming threat of drug-resistant infections.

## Introduction

The escalating threat of infections caused by drug-resistant bacteria poses a serious global public health challenge^1,2^. Projections estimate that annual deaths attributable to antibiotic resistance could reach 10 million by 2050^3^. However, the development of new antimicrobial agents has stagnated since the late 1990s, with approximately two-thirds of currently used antibiotics in clinical practice having been developed between the 1940s and 1960s^4,5^. Furthermore, only three new antibiotics with novel mechanisms of action have been developed in the six years leading up to 2023. This situation starkly illustrates the critical state of new antimicrobial development^6^.

Given this background, antimicrobial peptides (AMPs) have recently garnered attention as promising candidates for new antimicrobial agents^7–11^. AMPs are peptides involved in the innate immune response, acquired through evolution by a wide range of organisms, from prokaryotes to multicellular organisms. They are characterized by short amino acid sequences, typically 6-50 residues in length, and exhibit antimicrobial activity^9,12,13^. AMPs primarily exert their effects by destabilizing bacterial cell membranes and inducing lysis, a mechanism that is highly effective against drug-resistant bacteria. Moreover, experimental studies have demonstrated that AMPs are less prone to inducing resistance compared to conventional small-molecule antibiotics^8,12,14,15^. Recent discoveries of AMPs from the human gut microbiota^16^ and fragments of human protein sequences^17^ suggest that many AMPs remain undiscovered in nature. In this context, the discovery of novel AMPs is critically important for overcoming the limitations of existing antimicrobial agents. However, although over 3,000 AMPs have been identified to date^18^, only 17 peptide formulations have been approved by the Food and Drug Administration (FDA) for treating infectious diseases^19,20^, and only three of these target bacterial infections^21^. This indicates that challenges such as stability and toxicity pose significant barriers to the clinical application of AMPs^21^. Overcoming these obstacles requires establishing technologies capable of efficiently exploring and developing a more diverse array of AMPs than traditional experimental screening methods allow.

To address this, high-throughput computational screening has emerged as a promising approach. Previous studies have proposed classification methods based on the physicochemical properties of AMPs, as well as machine learning-based screening approaches^22^. More recently, deep learning-based classification models have been developed, and the application of artificial intelligence (AI) in the discovery of novel AMPs has gradually been explored^23,24^. Furthermore, protein language models (PLMs) have attracted attention as powerful tools for extracting structural and functional information from extensive protein datasets and have also been applied to the characterization of AMPs^25^.

Evolutionary Scale Modeling v2 (ESM-2)^26^ is a large-scale PLM trained on approximately 65 million proteins, which enables it to quantitatively capture the structural and functional features of proteins. Although there have been reports applying ESM-2 to AMP classification models^27,28^, previous studies have indicated that among the publicly available ESM-2 models, relatively smaller models have demonstrated superior performance^27–29^. This observation deviates from the common understanding that larger models with more parameters typically perform better, suggesting that a general-purpose language model, such as ESM-2, may not be fully optimized for specific tasks. These methods which use pre-trained PLMs directly do not fully harness its potential and are likely to be substantially improved by incorporating state-of-the-art natural language processing techniques. Furthermore, conventional approaches have employed artificially balanced datasets with a 1:1 ratio of positive to negative labels. Given that only a small fraction of naturally occurring peptides are AMPs, this ratio is inappropriate for models aimed at the exploration and development of novel AMPs. Due to these factors, although several computational approaches for AMP prediction have been proposed, few have led to the actual discovery of novel AMPs.

In this study, we developed a deep learning model aimed at discovering novel antimicrobial peptides (AMPs) to counter the threat posed by drug-resistant bacteria. Specifically, we introduce an AI model, AMP-Atlas (Fig. 1a), which utilizes ESM-2 to predict antimicrobial activity solely from peptide sequences and successfully identified novel AMPs. Unlike previous deep learning–based studies, AMP-Atlas combines Low-Rank Adaptation (LoRA)^30^, which efficiently adapts large-scale language models to specific tasks with limited data, with contrastive learning. This combination enables the efficient extraction of numerical representations of AMPs from ESM-2, even with scarce training data. Moreover, by employing contrastive learning, the differences in feature representations between AMPs and non-AMPs are accentuated, thereby enhancing the model’s ability to discriminate between AMPs. As a result, our proposed methodology outperforms conventional approaches and establishes a state-of-the-art model. Furthermore, using AMP-Atlas, we conducted a comprehensive search for AMPs in peptides and genomic sequences from various species and identified several novel AMP candidates that have not been previously reported (Fig. 1b). These findings demonstrate that AMP-Atlas is a highly promising foundational technology for the discovery of next-generation antimicrobial agents, especially for addressing the escalating problem of drug-resistant bacterial infections.

**Figure 1.**
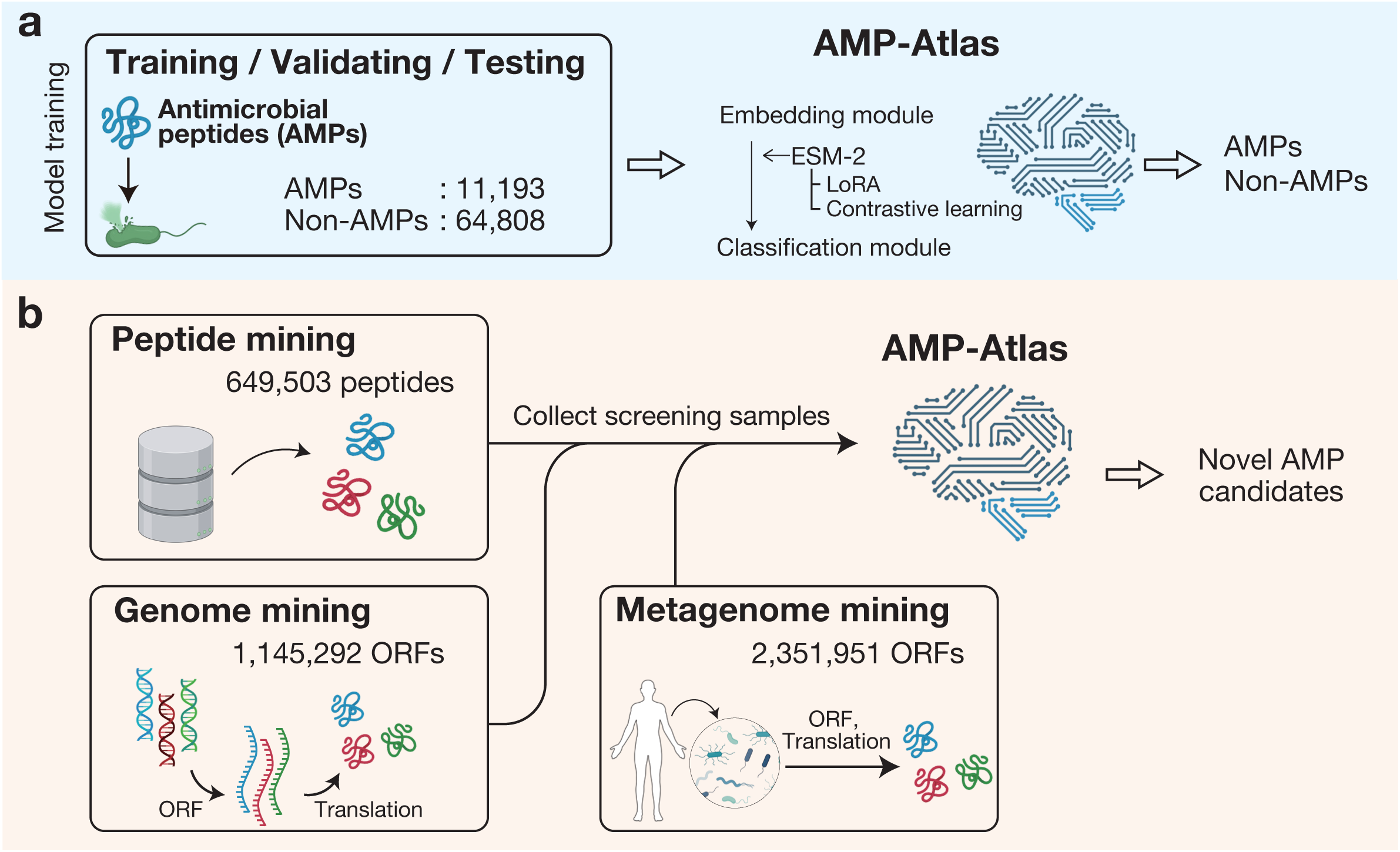
Overview of the construction and screening using AMP-Atlas. **(a)** AMP-Atlas was trained using peptide sequences labeled for antimicrobial activity. Amino acid sequences are input into Embedding Module for feature extraction. This module is based on ESM-2 and incorporates LoRA (Low-Rank Adaptation) and contrastive learning to acquire AMP-specific feature representations. Classification Module then predicts the presence or absence of antimicrobial activity based on the feature output of the Embedding Module, and the prediction is quantified as an AMP-likeness score. **(b)** Overview of screening for novel AMP candidates using AMP-Atlas. Novel AMP candidates are identified from existing peptide databases and genomic sequences using AMP-Atlas.

## Results

### AMP-Atlas: Construction of a deep learning framework for AMP prediction

To overcome the global challenges posed by the rise of drug-resistant bacteria and stagnation in novel antimicrobial drug development, we developed AMP-Atlas, a novel deep learning framework for accurately predicting antimicrobial peptides (AMPs) solely from sequence information. AMP-Atlas is built upon ESM-2, a state-of-the-art PLM, and achieves superior predictive performance compared to conventional methods by integrating Low-Rank Adaptation (LoRA)^30^ with contrastive learning (Fig. 1a).

The training dataset for AMP-Atlas was constructed using peptide sequences collected from multiple public databases, including APD^18^, dbAMPs^31^, DBAASP^32^, DRAMP^33^, SATPdb^34^, LAMP^35^, and CAMPR^36^, yielding 11,193 unique AMP sequences (Supplementary Fig. 1a, Materials and Methods). In addition, following previous studies^14,37^, 64,808 non-AMP peptide sequences (20–50 amino acids in length) were extracted from UniProt, excluding those annotated with “antimicrobial,” “antibacterial,” “antifungal,” or “antiviral.” The sequence length was restricted to 20-50 amino acids, consistent with the length distribution of reported AMPs (Supplementary Fig. 1b) and the availability of comprehensive screening methods for short peptides^14^. The non-AMP dataset comprising peptides with diverse functionalities other than antimicrobial activity, appropriately reflects the natural diversity of peptides and ensuring a training dataset well-suited for AMP discovery. A comparison of the amino acid compositions of the collected AMPs and non-AMPs revealed that AMPs have higher proportions of cationic amino acids, such as lysine and histidine, as well as the hydrophobic amino acid glycine, which is consistent with previously reported characteristics^8,38,39^ (Supplementary Fig. 1c). To prevent overfitting, the entire dataset was randomly partitioned into training, validation, and test sets in a ratio of 8:1:1.

Next, we examined the model architecture of AMP-Atlas. To obtain numerical representations of three-dimensional structures and physicochemical properties from amino acid sequences, we focused on ESM-2 and leveraged its peptide sequence embeddings. The AMP-Atlas architecture consists of an Embedding Module, responsible for extracting sequence features, and a Classification module that determines the presence or absence of antimicrobial activity (Fig. 2a). However, because the feature representations generated by ESM-2 are universal protein sequence embeddings, they may not sufficiently capture the AMP-specific characteristics. To enhance AMP discrimination, we posited that additional domain-specific fine-tuning was necessary. Specifically, we hypothesized that introducing Low-Rank Adaptation (LoRA) ^30,40,41^, a method that adapts general-purpose large language models to specific tasks by approximating a subset of model parameters with low-rank matrices, would improve the classification accuracy between AMPs and non-AMPs. LoRA enables efficient fine-tuning, which is particularly advantageous in life sciences, where available data are often limited. Furthermore, to strengthen the extraction of AMPs representations from ESM-2, we integrated the concept of contrastive learning into the model. Contrastive learning enhances separability in the feature space by minimizing the distance between samples with the same label (AMP or non-AMP) and maximizing the distance between samples with different labels. Specifically, we applied a multi-class N-pair loss^42^ constraint to the numerical vectors obtained via ESM-2 and LoRA, aiming to yield embeddings that reflect label similarities (Materials and Methods).

**Figure 2.**
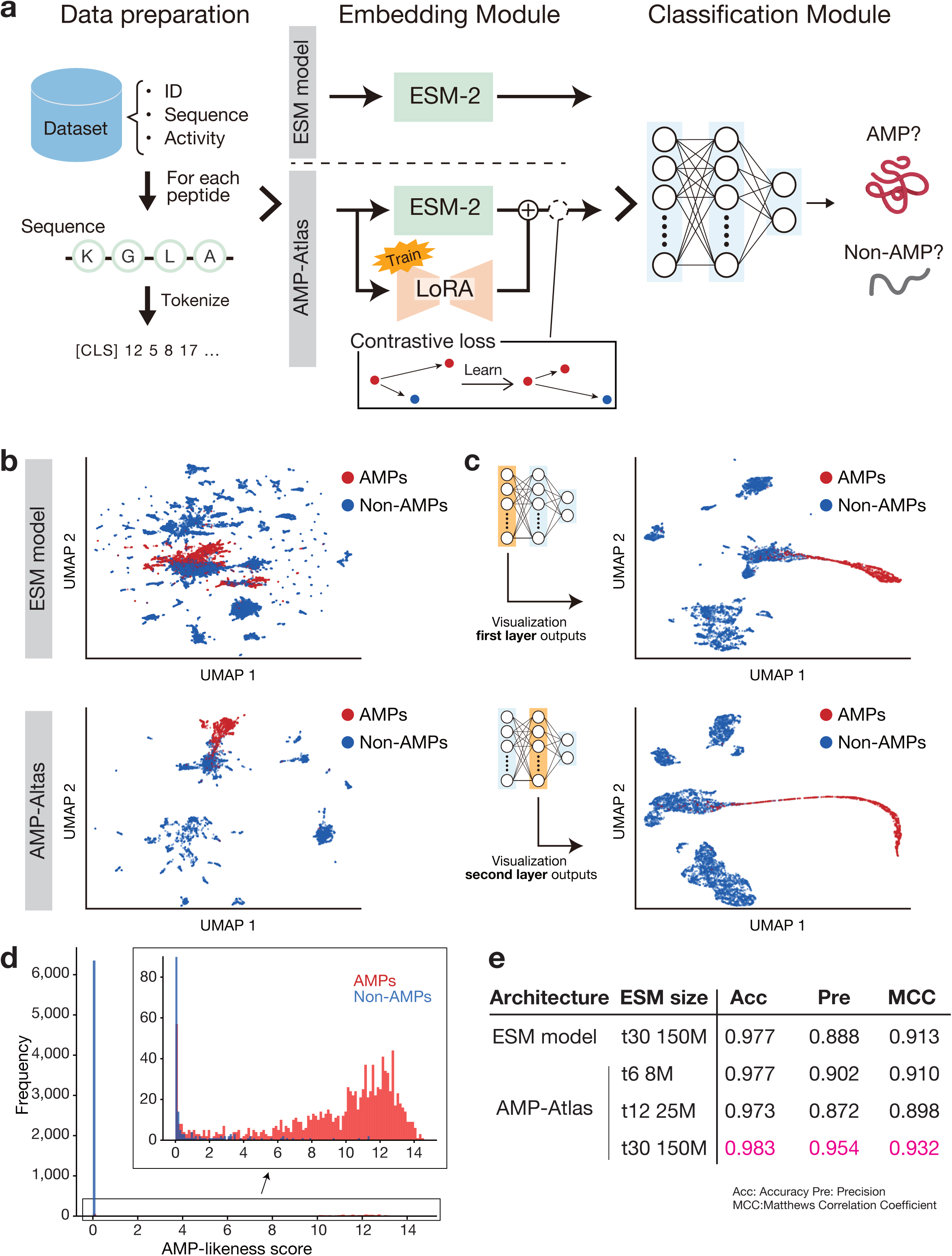
AMP-Atlas efficiently extracts and learns AMPs features. **(a)** Data collection and model architecture. Training data collected from public databases included IDs, amino acid sequences, and information regarding antimicrobial activity. Two versions of Embedding Module were compared: one based solely on ESM-2 (upper panel), and the AMP-Atlas version incorporating LoRA and contrastive learning (lower panel). The Classification Module predicts antimicrobial activity based on the features extracted by the Embedding Module. **(b)** Two-dimensional visualization of peptide numerical representations from the test data generated by Embedding Module. Dimensionality reduction was performed using UMAP. The upper panel shows the results for the ESM-2 only model, and the lower panel shows those for AMP-Atlas (ESM-2 + LoRA + contrastive learning). Red dots represent AMPs, and blue dots represent non-AMPs. **(c)** Two-dimensional plots of the latent space across each layer of the Classification Module (upper panel: first layer; lower panel: second layer). Dimensionality reduction was performed using UMAP. Red dots indicate AMPs, and blue dots indicate non-AMPs. **(d)** Distribution of AMP-likeness scores in the test data. The horizontal axis represents the AMP-likeness score, and the vertical axis represents the frequency. The inset graph shows an enlarged view of the data, with frequency values ranging from 0 to 90. **(e)** Comparison of model performance as a function of network architecture and ESM-2 model size. The highest values of each evaluation metric are highlighted in pink. Metrics included accuracy (Acc), precision (Pre), and Matthews Correlation Coefficient (MCC).

### Enhanced predictive performance via LoRA and contrastive learning

To evaluate the effectiveness of our proposed method, we constructed two models: (1) a model using only ESM-2 and (2) a model incorporating both LoRA and contrastive learning with ESM-2 (AMP-Atlas). We compared their performance using the same test dataset (Fig. 2a). The results demonstrate that AMP-Atlas outperformed the ESM-2-only model across most performance metrics (Table 1). Notably, the substantial improvement in precision, a key metric for AMP discovery, suggests that our approach effectively reduces false positives, thereby enabling more efficient AMP discovery.

**Table 1.**
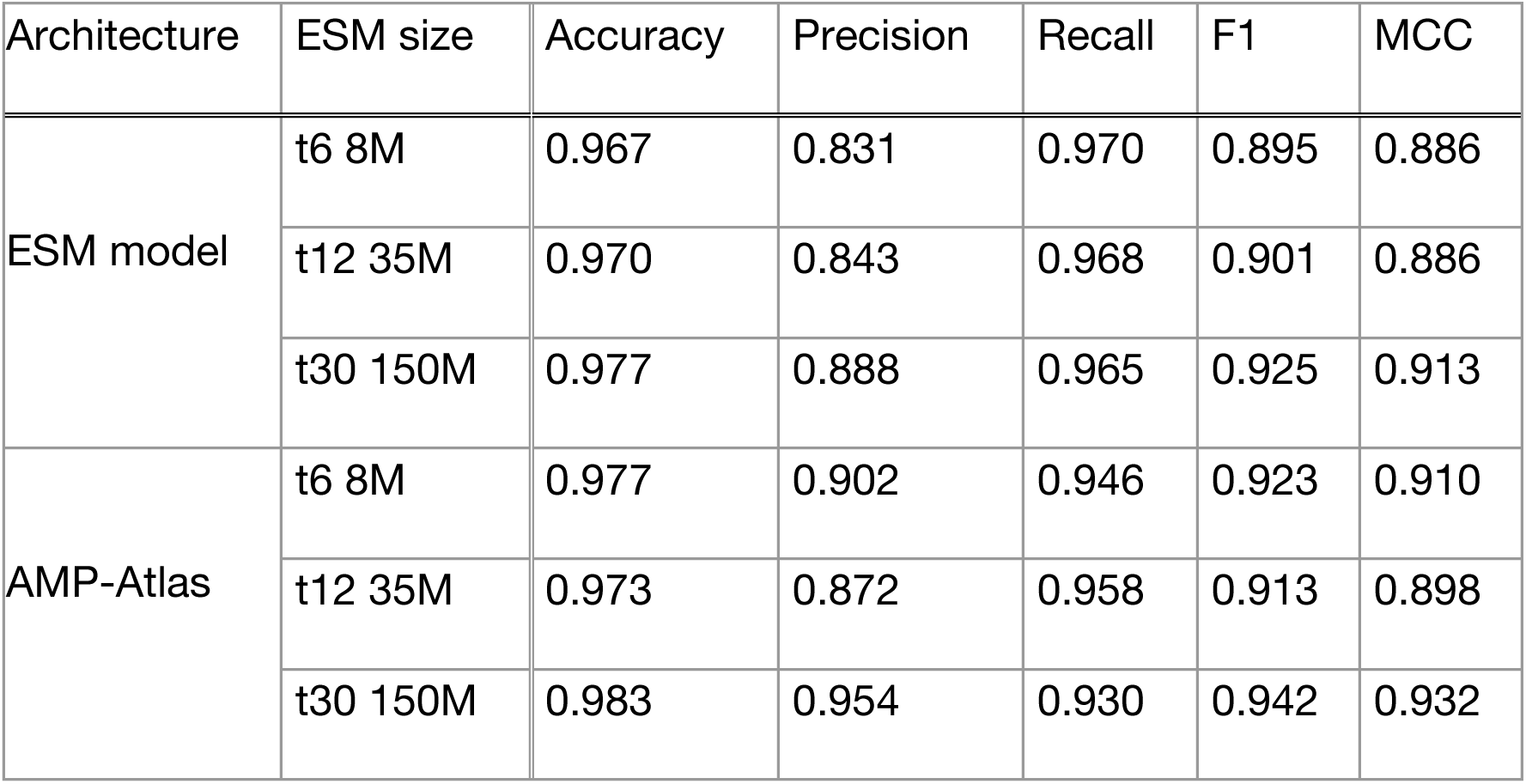
The ablation study of Embedding Module.

Furthermore, we visualized the distribution of the feature vectors produced by the Embedding Module of each model in two dimensions using UMAP (uniform manifold approximation and projection)^43^. Compared to the ESM-2-only model, AMP-Atlas exhibited a more distinct separation between AMPs and non-AMPs (Fig. 2b), indicating that the introduction of LoRA and contrastive learning effectively captured AMP-specific features. In addition, visualization of the latent space across the layers of the Classification module revealed that the separation between AMPs and non-AMPs became increasingly pronounced as the representations progressed toward the output layer (Fig. 2c). This finding suggests that the multilayer structure progressively extracts higher-order features of AMPs. Collectively, these results demonstrate that AMP-Atlas, incorporating both LoRA and contrastive learning, not only enhances the accuracy of AMP discrimination but also yields more separable numerical representations, thereby validating the effectiveness of these techniques.

### Comparison of AMP-Atlas’s predictive performance with existing methods

AMP-Atlas quantifies the antimicrobial activity of an input peptide sequence as an AMP-likeness score (Fig. 2a), which reflects the likelihood that the peptide exhibits antimicrobial properties. Visualization of the AMP-likeness score distribution in the test data revealed that non-AMP peptides clustered near a score of 0, whereas AMPs tended to have higher scores (Fig. 2d). This suggests that our approach may enable high-precision identification of the relatively scarce AMPs present in nature. Based on these findings, we set a threshold such that peptides with an AMP-likeness score of 4 or higher were considered to possess antimicrobial activity.

Next, we compared the predictive performance of AMP-Atlas with that of existing AMP prediction methods. For comparison, we selected two state-of-the-art deep learning–based approaches, Diff-AMP^28^ and Cao et al.^44^, and we retrained these methods on our dataset before evaluating their performance on the same test data (Materials and Methods). The results demonstrate that AMP-Atlas is capable of identifying AMPs with higher accuracy and precision than existing methods (Table 2).

**Table 2.** Comparison of the performance of AMP-Atlas with previous studies.

| Method | Accuracy | Precision | Recall | F1 score | MCC |
| --- | --- | --- | --- | --- | --- |
| Diff-AMP<sup>28</sup><br>(re-trained) | 0.978 | 0.931 | 0.920 | 0.926 | 0.913 |
| Cao et al.<sup>44</sup><br>(re-trained) | 0.972 | 0.935 | 0.876 | 0.904 | 0.889 |
| AMP-Atlas<br>(Our model) | 0.983 | 0.954 | 0.930 | 0.941 | 0.932 |

Furthermore, our experiments provide new insights into the performance of applying LoRA to PLMs. Conventionally, when using ESM-2 for AMP prediction, it has been suggested that larger models with more parameters are not necessarily optimal^27–29^. However, in this study, by combining LoRA with contrastive learning on ESM-2, we observed that the classification performance for AMPs improved as the model size (i.e., the number of parameters) increased (Fig. 2e, Table 1). Notably, there was a significant improvement in precision, which is a critical metric for AMP discovery. This finding indicates that LoRA and contrastive learning can fully harness the latent capabilities of large-scale PLMs and enhance their adaptation to specific tasks, providing important insights into the development of more effective AMP prediction models.

### Prediction of novel AMPs by AMP-Atlas

AMPs are believed to have evolved in various organisms as a defense mechanism against bacterial threats, and it is anticipated that many undiscovered AMPs exist in nature^12^. Leveraging the high predictive performance of AMP-Atlas, we searched for novel AMPs using existing peptide databases and genomic sequences (Fig. 3a). We performed peptide mining on 649,503 eukaryotic peptides (20–50 amino acids in length) retrieved from the UniProt database (Fig. 3b) and genome mining on 1,145,292 open reading frames (ORFs) predicted from the genomic sequences of model organisms, including humans (Fig. 3c) (Materials and Methods). As a result, AMP-Atlas detected peptides with high AMP-likeness scores (>4) in both datasets (Fig. 3d). Interestingly, the proportion of high-scoring peptides was very low (1.2% for peptide mining and 0.2% for genome mining), suggesting that these values reflect a naturally low prevalence of AMPs. Moreover, because AMPs interact with negatively charged bacterial cell membranes, they are known to exhibit a net positive charge^9,45^. Consistent with this, the AMP candidate peptides predicted by AMP-Atlas demonstrated a net positive charge compared to the non-AMP candidate peptides (Fig. 3e). This finding suggests that AMP-Atlas effectively captured the physicochemical properties of AMPs during the training process.

**Figure 3.**
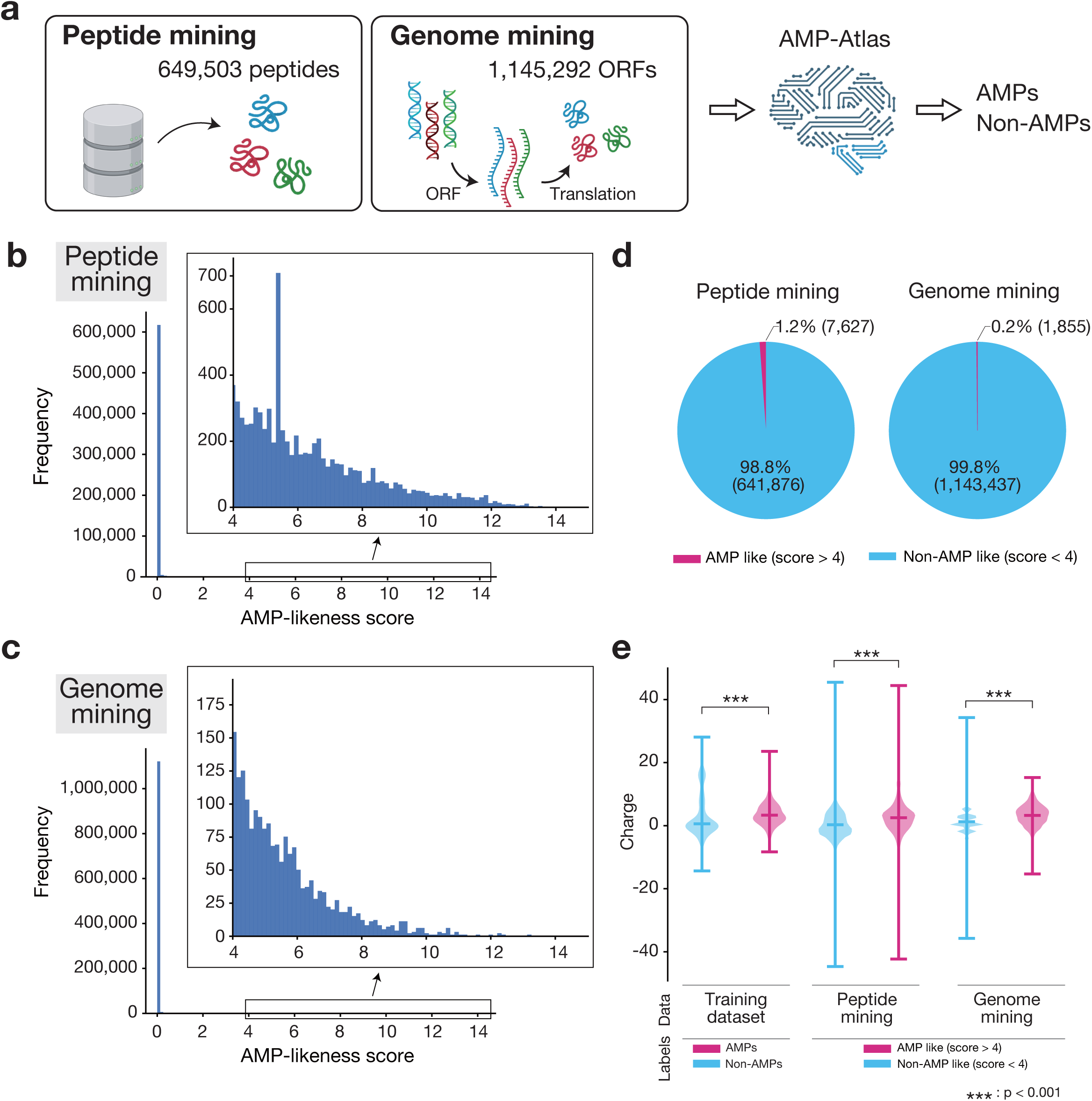
AMP-Atlas discovers novel AMP candidates from peptide and genomic data. **(a)** Overview of peptide mining and genome mining. Screening was performed using AMP-Atlas on eukaryotic peptides obtained from the UniProt database and on ORFs predicted from the genomic sequences of model organisms. The numbers indicate the sequence counts used for screening. **(b, c)** Results of peptide mining (b) and genome mining (c). The horizontal axis represents the AMP-likeness score, and the vertical axis represents frequency. The inset graphs show an enlarged view of the data with AMP-likeness scores of 4 or higher. **(d)** The proportion of AMP candidate peptides predicted by peptide mining (left) and genome mining (right). Red indicates AMP candidate peptides (AMP-likeness score > 4), and blue indicates other peptides (AMP-likeness score < 4). **(e)** Distribution of the net charge of peptide sequences across the datasets. The horizontal axis represents the dataset (left: training data; center: peptide mining results; right: genome mining results), and the vertical axis represents the net charge. ***: p < 0.001 (Welch’s *t*-test).

### Structural validity of AMP-Atlas predictions for antimicrobial peptides

AMPs are widely distributed across diverse species and are known to exhibit characteristic three-dimensional structures and surface property patterns associated with antimicrobial activity^9,12,15^. Therefore, among the AMP candidates identified via peptide mining (Fig. 3b), we analyzed the species of origin, three-dimensional structures, and surface charges of peptides with high AMP-likeness scores (Fig. 4a). AMP-Atlas predicted AMP candidates from a variety of sources, including plants, insects, amphibians, fish, annelids, and mammals. Some candidates, such as peptides derived from Jute, Adineta, and Rainbow trout, represent functionally uncharacterized sequences not registered in existing AMP databases.

**Figure 4.**
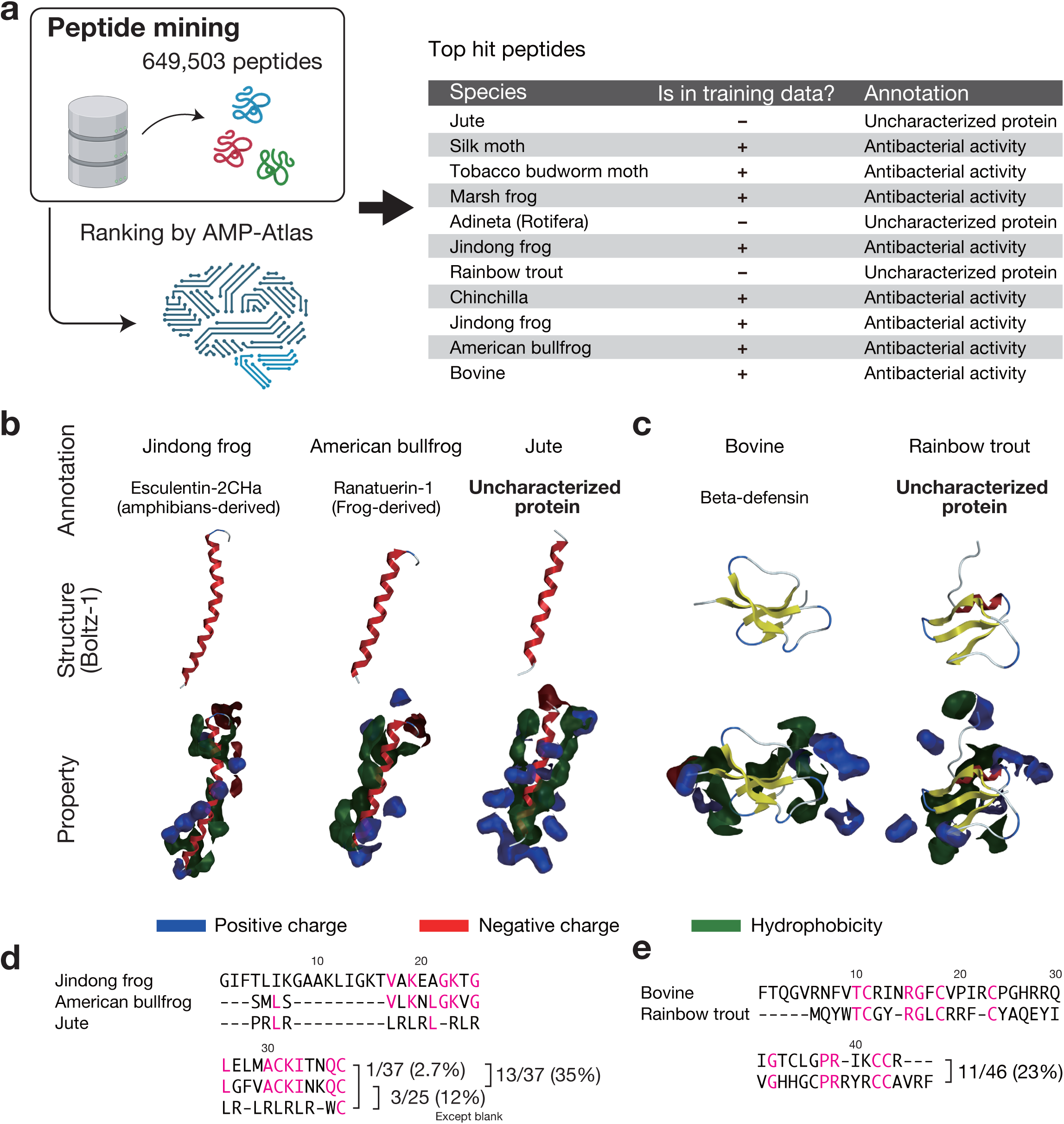
AMP candidates exhibit structural properties indicative of antimicrobial activity. **(a)** Information on peptides with high AMP-likeness scores identified via peptide mining. For the top-scoring peptides obtained by AMP-Atlas, the table shows the species of origin, inclusion in the training data, and their annotations in UniProt. **(b, c)** Three-dimensional structures (upper panel) and surface charge distributions (lower panel) of peptides with high AMP-likeness scores. Panel (b) shows an example of a peptide with a helical structure, whereas panel (c) shows an example of a peptide with a β-sheet structure. The highlighted structures are functionally uncharacterized peptides. The three-dimensional structures were predicted using Boltz-1, and the surface charge distributions were calculated using MOE (blue: positively charged regions, red: negatively charged regions, green: hydrophobic regions). **(d, e)** Amino acid sequences of peptides with high AMP-likeness scores. Panel (d) shows an example of a peptide with a helical structure, whereas panel (e) shows an example of a peptide with a β-sheet structure. Highlighted amino acid residues are conserved across peptide. The numbers indicate the degree of sequence similarity.

The predicted three-dimensional structures of these AMP candidates were modeled using Boltz-1^46^, revealing that most adopted amphipathic α-helical structures (Fig. 4b) or β-sheet structures (Fig. 4c). These conformations are typical of known AMPs and are related to their mechanisms of action on bacterial cell membranes^8,15,45,47^. Additionally, analysis of the surface charge distribution with MOE showed that these peptides exhibited clearly segregated regions of positive charge and hydrophobicity, further confirming their amphipathic nature (Materials and Methods). Based on the above findings, these functionally uncharacterized peptides predicted by AMP-Atlas to be AMPs exhibited properties similar to those of known AMPs. These observations suggested that they are likely to possess antimicrobial activity. Importantly, the sequence identity between these AMP candidates and known AMPs was extremely low (ranging from 2.7% to 23%, Fig. 4d and Fig. 4e). These findings suggest that AMP-Atlas captures higher-order features rather than just predictions based on sequence similarities.

### Metagenome mining for the discovery of novel AMP candidates

Diverse microbial communities in nature have evolved AMPs as a means of attack and defense during competitive and symbiotic interactions. In particular, microbiomes, where various bacterial species co-occur, have been reported to harbor community-specific AMPs^48^. Uncultured microorganisms, which cannot be accessed via conventional cultivation methods, are considered promising sources of novel AMPs. Therefore, we attempted to identify novel AMPs using environmental metagenomic data.

Commensal microbiota coexist with their human hosts. Consequently, substances produced by these bacteria are expected to have minimal adverse effects on human tissues^49,50^. Based on this hypothesis, we aimed to discover AMPs with high safety profiles from the metagenomes of the commensal microbiota, specifically the gut and oral microbiomes (Fig. 5). We obtained metagenomic data for the human gut microbiome (Study ID: MGYS00005762) and the oral microbiome (Study ID: MGYS00006562) from the MGnify^51^ database (Fig. 5a). ORFs ranging from 20 to 50 amino acids in length were predicted using Prodigal^52^, and these ORFs were subsequently screened using AMP-Atlas. As a result, 332,874 ORFs were predicted from the gut microbiome and 2,019,077 were predicted from the oral microbiome, yielding a combined total of 46,999 candidate AMP sequences (Fig. 5b). The metagenome-derived candidate AMP sequences exhibited a net positive charge, similar to that observed for known AMPs (Fig. 5c). This finding is consistent with the results shown in Fig. 3e, suggesting that the candidate AMP sequences predicted from the metagenomes likely possess antimicrobial activity. Furthermore, when the three-dimensional structures of peptides with high AMP-likeness scores were predicted, many were found to adopt α-helical conformations, and their surface charge distributions demonstrated amphipathicity (Fig. 5d).

**Figure 5.**
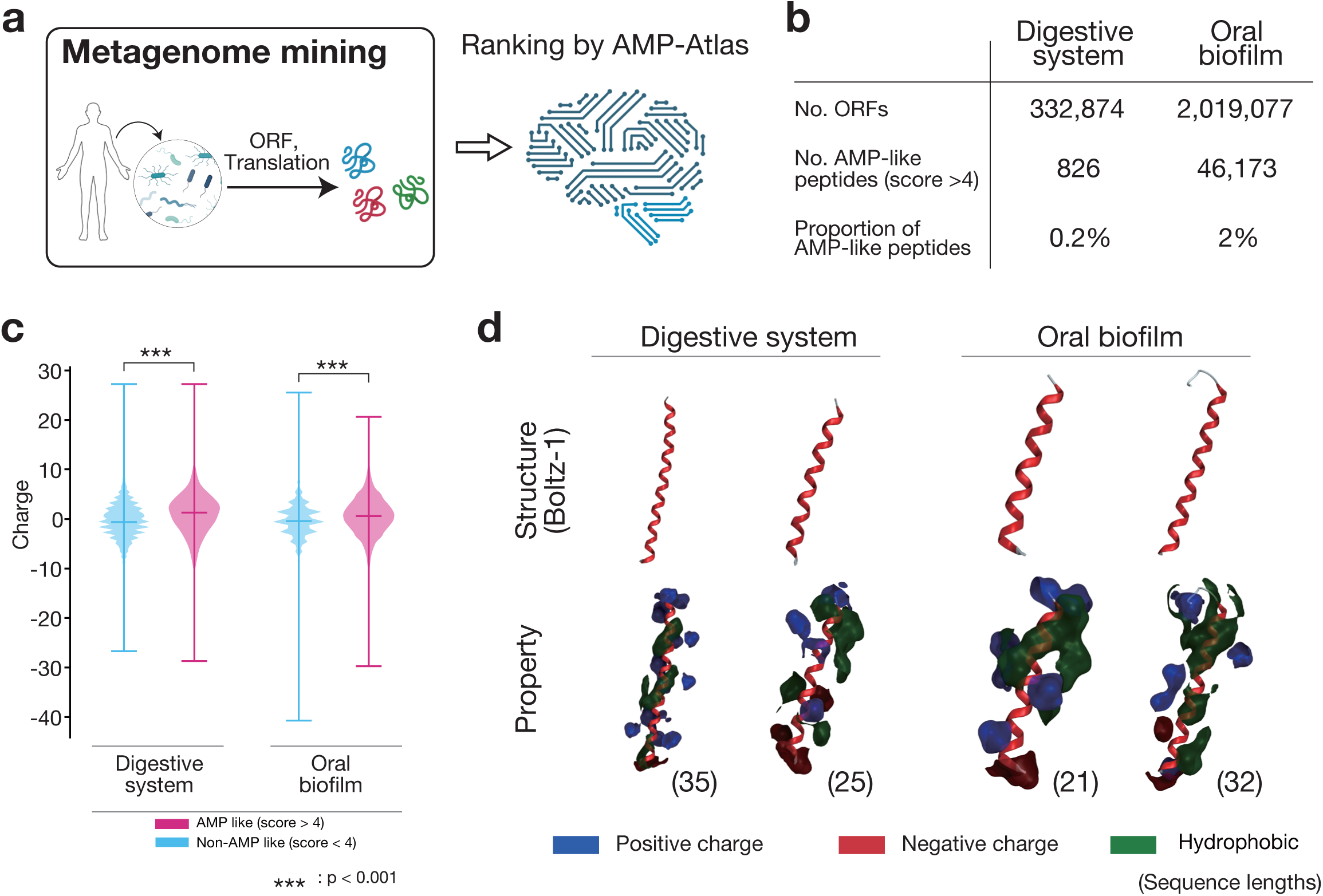
The human commensal microbiome metagenome contains a vast number of AMP candidates. **(a)** Overview of metagenome mining. ORFs were predicted from metagenomic data obtained from the human commensal microbiota (gut and oral microbiomes), translated into amino acid sequences, and subsequently screened using AMP-Atlas. **(b)** Results of the AMP-Atlas screening. Peptides with an AMP-likeness score of 4 or higher were designated as AMP candidates. **(c)** Distribution of the net charge of peptide sequences in each dataset. The horizontal axis represents the dataset (left: gut microbiome, right: oral microbiome), and the vertical axis represents the net charge. ***: p < 0.001 (Welch’s *t*-test). **(d)** Three-dimensional structures (upper panel) and surface charge distributions (lower panel) of peptides with high AMP-likeness scores. The three-dimensional structures were predicted using Boltz-1, and the surface charge distributions were calculated using MOE (blue: positively charged regions, red: negatively charged regions, green: hydrophobic regions). The numbers indicate the peptide sequence lengths.

Collectively, these results indicate that AMP-Atlas can efficiently identify structurally and functionally valid AMP candidates from metagenomic data.

## Discussion

Antimicrobial resistance poses a serious global health challenge, and its threat continues to escalate. The emergence of resistant pathogens against existing antibiotics, coupled with the stagnation of novel antimicrobial drug development, necessitates the urgent establishment of new therapeutic strategies. To address this issue, we developed AMP-Atlas, an AI-based system for the discovery of antimicrobial peptides (AMPs). AMP-Atlas is built on the protein language model ESM-2 and combines LoRA with contrastive learning to achieve a prediction accuracy that surpasses existing AMPs prediction methods. Furthermore, using AMP-Atlas, we conducted large-scale mining of peptide and genomic data to identify novel AMP candidates. In addition, we identified a vast number of AMP candidates from the metagenomic data of the human commensal microbiota, demonstrating the potential of metagenomes as an untapped resource for drug discovery. These findings underscore the potential of AI-driven approaches for the exploration and generation of AMPs and provide a new strategy for combating drug-resistant pathogens.

AMP-Atlas’s technical innovation lies in its specialization of ESM-2, originally designed to acquire universal protein sequence embeddings, for the prediction of function-specific AMPs. Traditionally, large-scale PLMs such as ESM-2, despite their versatility, have sometimes fallen short in performance on specific tasks. To overcome this limitation, we employed two strategies: (1) efficient fine-tuning via LoRA and (2) feature optimization through contrastive learning. LoRA approximates a subset of the model’s parameters with low-rank matrices, thereby significantly reducing the number of training parameters while enabling efficient fine-tuning^30^. This approach makes it possible to adapt the ESM-2 for AMPs prediction, even in computationally constrained environments. Contrastive learning optimizes the arrangement of samples in the feature space based on label information by drawing together samples with the same label (AMPs or non-AMPs) and separating those with different labels. In this study, we employed Multi-class N-pair Loss^42^ to acquire feature representations that allow for clear separation between AMPs and non-AMPs. By integrating these techniques, AMP-Atlas demonstrated significantly higher performance across all evaluation metrics (Accuracy, Precision, Recall, F1 score, and MCC) compared to existing AMP prediction methods^28,44^ (Table 1). Notably, the marked improvement in precision suggests that AMP-Atlas effectively minimizes false positives, thereby enabling more efficient AMP discovery. This finding has the potential to substantially reduce experimental validation costs and represents an important contribution to accelerating AMP drug discovery research.

The utility of AMP-Atlas was demonstrated by its ability to identify novel AMP candidates from large-scale peptide and genomic sequences (Fig. 3), as well as by its discovery of AMP candidates from metagenomic data (Fig. 5). The proportion of peptides predicted to possess antimicrobial activity through AMP-Atlas screening ranged from only 0.2% to 2% of the total, which is consistent with the fact that only a very small fraction of peptides in nature exhibit antimicrobial activity. This finding indicates that AMP-Atlas can efficiently detect this rare subset of AMPs. Moreover, it has been reported that commensal bacteria, through their co-evolution with their human hosts, produce proteins that are less likely to trigger adverse events, such as inflammation, or exhibit high immunogenicity compared to those produced by environmental bacteria^49,50^. Therefore, AMP candidate peptides identified from the human commensal microbiota using AMP-Atlas are expected to possess similar safety characteristics and hold significant potential for advancing the clinical application of AMPs.

Nevertheless, this study has several limitations. First, the current version of AMP-Atlas is trained based on AMPs information registered in multiple existing databases. However, these databases may be biased toward AMPs from specific species or belonging to certain structural classes. A future challenge is to collect a more diverse range of AMP data and expand the training dataset. Second, the current AMP-Atlas is designed to predict peptides comprising 20–50 amino acids. Although this range is justified by the reported comprehensive screening method for short peptides^14^ and by the length distribution of the reported AMPs (Supplementary Fig. 1b), naturally occurring AMPs can be shorter or longer. In the future, it will be necessary to extend the range of peptide lengths considered for prediction to discover a more diverse array of AMPs. Third, although AMP-Atlas predicted a large number of AMP candidates, these results represent a hypothesis that requires experimental validation. *In vitro* and *in vivo* biological evaluations are needed to elucidate the antimicrobial activity, cytotoxicity, and mechanism of action of the candidate peptides. By incorporating such experimental data as feedback into AMP-Atlas, it is expected that the performance of the model can be further improved, enabling more accurate functional predictions and establishing AMP-Atlas as a foundational platform for peptide drug discovery.

Our AMP-Atlas could serve as a game-changer in the battle against drug-resistant bacteria. AMP-Atlas possesses the ability to efficiently mine unknown AMPs present in nature and generate a vast array of AMP candidates from diverse data sources. This represents a crucial breakthrough in the era when existing antibiotics are losing their efficacy. Moreover, AMP-Atlas holds the potential to accelerate next-generation antimicrobial drug development, including *de novo* AMP design and the design of AMPs targeted at specific pathogens. This study provides a foundational technology to confront the greatest threat to humanity, antimicrobial resistance, and paves the way for a new era of AMP drug discovery.

## Materials and Methods

### Training datasets

In this study, sequence data for AMPs and Non-AMPs were collected from multiple public databases. AMP data were obtained from the following AMP-specific public databases: dbAMPs^12^, APD^18^, DBAASP^32^, DRAMP^33^, SATPdb^34^, LAMP^35^ and CAMPR^36^ (downloaded on July 27, 2023). For Non-AMPs, sequences were downloaded from the protein database UniProt, excluding those containing any of the annotations “antimicrobial,” “antibacterial,” “antifungal,” or “antiviral,” and selecting proteins with an annotation score of at least 2. The sequence lengths were restricted to the range of 20–50 amino acids. This criterion was based on the length distribution of reported AMPs (Supplementary Fig. 1b) and the existence of comprehensive screening methods for short peptides^14^. In the downloaded amino acid sequences, J (ambiguous residue that cannot be assigned as either isoleucine or leucine) and non-standard amino acids were replaced with X (totally unknown). Additionally, peptides with branched backbone structures were excluded from analysis. Finally, duplicate sequences were removed, resulting in 11,193 unique AMPs sequences and 64,808 Non-AMP sequences used for training and analysis. To prevent overfitting and evaluate the generalization performance of the model, the dataset was randomly split into training, validation, and test sets at an 8:1:1 ratio. The test data were reserved solely for the final performance evaluation and were not used during model training.

### Architecture of the Embedding Module

AMP-Atlas is a deep learning model composed of Embedding Module, which is responsible for extracting sequence features, and Classification Module, which predicts the presence or absence of antimicrobial activity (Fig. 2a). Feature extraction from the protein sequences was performed using ESM-2^26^. To specialize the general-purpose protein embeddings generated by ESM-2 for extracting features related to antimicrobial activity, LoRA^30^ and contrastive learning were incorporated. LoRA enables efficient adaptation to a specific task by adding low-rank matrix adapters to certain substructures of a large language model (LLM). This is formulated as:

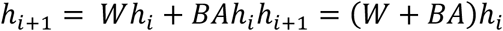

where *h_i_* and *h_i_*_+2_ are the input and output vectors, respectively; *A* is a matrix of dimensions (*h_i_*, *r*) and *B* is a matrix of dimensions (*h_i_*_+1_, *r*). *W* represents the fixed weights of the corresponding substructure block of the pre-trained LLM. The Hugging-Face PEFT (Parameter-Efficient Fine-Tuning) library (version 4.33.3) was used to implement LoRA. The TaskType for LoRA was specified as “FEATURE_EXTRACTION,” and LoRA was applied to the transformation layers (“query,” “key,” “value”) within the attention structure of ESM-2. The LoRA hyperparameters (r = 8, lora_alpha = 15.48) were determined using Bayesian optimization with Optuna^53^.

Contrastive learning is a training strategy that optimizes the arrangement of samples in the feature space based on label information by bringing closer samples with the same label (AMPs or Non-AMPs) and pushing apart those with different labels. In this study, multi-class N-pair loss^42^ was used as the contrastive loss function:

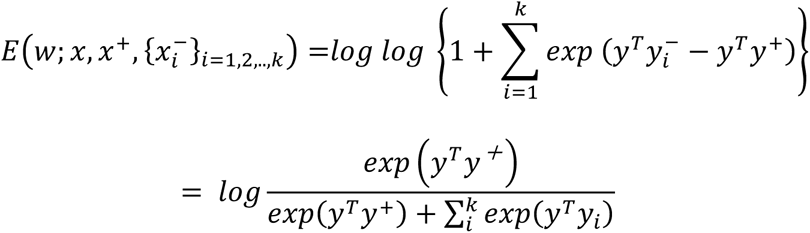

Here, *x* represents an anchor peptide sequence, *x*^+^ denotes a peptide sequence with a positive label (AMPs), and *x*^−^ denotes a peptide sequence with a negative label (Non-AMPs). *y*, *y*^+^, and *y*^−^ correspond to the embedding vectors of the anchor, positive, and negative peptides, respectively. Variable *w* represents the parameters of the Embedding Model (ESM-2 and LoRA).

### Architecture of the Classification Module

Classification Module takes the peptide feature vectors extracted by the Embedding Module as input and outputs a prediction of antimicrobial activity (AMPs or Non-AMPs) as output. This module is composed of a multilayer perceptron (MLP) with hidden layer sizes of [128, 64, 2]. The number of units in each layer was determined by Bayesian optimization using Optuna^53^.

### Training of AMP-Atlas

AMP-Atlas was trained by jointly optimizing the Embedding and Classification Modules. The AdamW optimizer^54^ was used with a weight decay of 0.044 and a learning rate of 0.00057; these hyperparameters were determined using Optuna^49^. A batch size of 32 was used, and 80% of the training dataset was randomly selected for model training. The loss function is defined as the sum of the cross-entropy loss and contrastive losses.

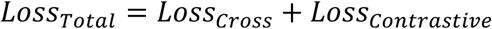

where *Loss_cross_* is the cross-entropy loss (CEL), defined as:

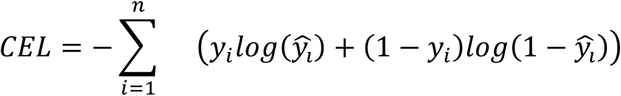

Here, *y_i_* and *ŷ_i_* denote the true label and the predicted label for the *i* -th sample, respectively. Early stopping was implemented with a patience of 10 epochs, meaning that training was halted if the loss in the validation dataset did not improve for 10 consecutive epochs.

### Performance evaluation metrics

The predictive performance of AMP-Atlas was evaluated using five metrics: accuracy (ACC), precision (PRE), recall (REC), F-score (FSC), and Matthew’s correlation coefficient (MCC). These metrics are defined as follows:

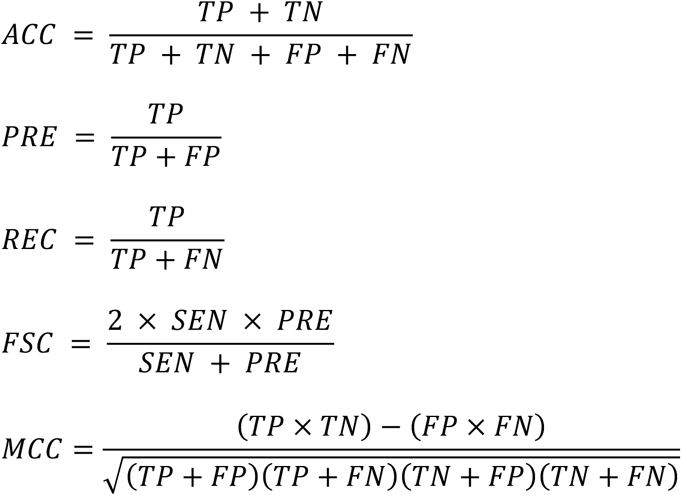

where TP, TN, FP, and FN denote the true positives, true negatives, false positives, and false negatives, respectively.

### Comparison with existing methods

To evaluate the performance of AMP-Atlas, a comparison was made with existing state-of-the-art AMP prediction methods (Diff-AMP^28^, Cao et al.^44^). These methods were retrained using a comprehensive dataset constructed from multiple databases used in this study, and their performances were evaluated using the same test data to ensure fairness.

### Discovery of novel AMP candidates from large-scale sequence data

The pre-trained AMP-Atlas was used to discover novel AMP candidates from large-scale sequence data. For peptide mining (Fig. 3b), eukaryotic peptide sequences with lengths of 20–50 amino acids were obtained from UniProt (downloaded on December 14, 2023). After excluding peptides containing non-standard amino acids, a total of 649,503 peptides were screened using AMP-Atlas. For genome mining (Fig. 3c), open reading frames (ORFs) were obtained from OpenProt^55^, consisting of 60–150 bp sequences derived from *Homo sapiens*, *Drosophila melanogaster*, *Mus musculus*, *Rattus norvegicus*, *Caenorhabditis elegans*, *Saccharomyces cerevisiae*, and *Danio rerio*, totaling 1,145,292 sequences (downloaded on October 12, 2024). During screening, the predicted amino acid sequences translated from the genomic data were used.

### Metagenome mining

To leverage genetic information from uncultured microorganisms, AMP candidates were also explored using metagenomic data (Fig. 5). Metagenomic data (FASTA format) from the human gut microbiome (Study ID: MGYS00005762) and the oral microbiome (Study ID: MGYS00006562) were downloaded from the MGnify^51^ database (downloaded on February 10, 2025). ORFs with lengths between 20 and 50 amino acids were predicted from these data using Prodigal^52^ (version 2.6.3) with the “-p meta” option, and the predicted ORF sequences were screened using AMP-Atlas to identify AMP candidate sequences.

### Calculation of net charge of peptides

The net charge (calculated at pH 7.4) of peptides in the training, peptide mining, genome mining, and metagenome mining datasets was computed using the Bio.SeqUtils.ProtParam.ProteinAnalysis function from the BioPython package (version 1.84).

### Structural analysis of AMP candidate peptides

The three-dimensional structures of AMP candidate peptides predicted by AMP-Atlas were determined using Boltz-1^46^. Multiple sequence alignments (MSAs) required for structure prediction were generated using mmseqs2 server^56^.

Surface charge and hydrophobic patch analyses were performed using the Molecular Operating Environment (MOE; Chemical Computing Group Inc.). Prior to the analysis, hydrogen atoms were added, and structural optimization was performed using the Amber10: Extended Hückel Theory (EHT) ^57^ force field. The Protein Patch module in MOE was used to quantitatively calculate the interaction regions on the protein surface, specifically electrostatic patches and hydrophobic patches. The charge map was generated by partitioning the surface based on local electric fields obtained via Ewald summation^58^, with regions exceeding 40 kcal/mol/C in absolute value defined as electrostatic patches^59,60^. Regions with an area of 40 Å² or less were excluded. Hydrophobic regions were determined using an octanol-water partition coefficient (logP)-based hydrophobic potential map, with regions having a potential above 0.09 kcal/mol defined as hydrophobic patches^60^; regions with an area of 50 Å² or less were excluded.

### Statistical Analysis

Statistical analyses (Fig. 3e, Fig. 5c) were performed using the Python package SciPy (version 1.13.0). As a preprocessing step, the most extreme 10% of the data for each dataset were excluded. Welch’s *t*-test was used for statistical comparisons, and a one-sided test was adopted with a significance level of *p* < 0.05.

## ACKNOWLEDGEMENTS

This work was supported by JST, PRESTO Grant Number JPMJPR22R6, Japan to H.S. This work was also partly supported by KAKENHI grants from the Japan Society for the Promotion of Science (JSPS) to H.S. (24H01755 and 23K18502), as well as Takeda Science Foundation. We thank H. Aso, S. Ohno, and other laboratory members for discussion and K. Tanaka for help with preparation of the manuscript.

## AUTHOR CONTRIBUTIONS

H. S. conceived of, designed, and supervised the study. Y.O. developed AMP-Atlas and, with inputs from D.K. and Y.W., performed all computational analyses. Y.O. and H.S. jointly wrote the manuscript with the comments from all authors.

## Supplementary Information

**Supplementary Figure 1.**
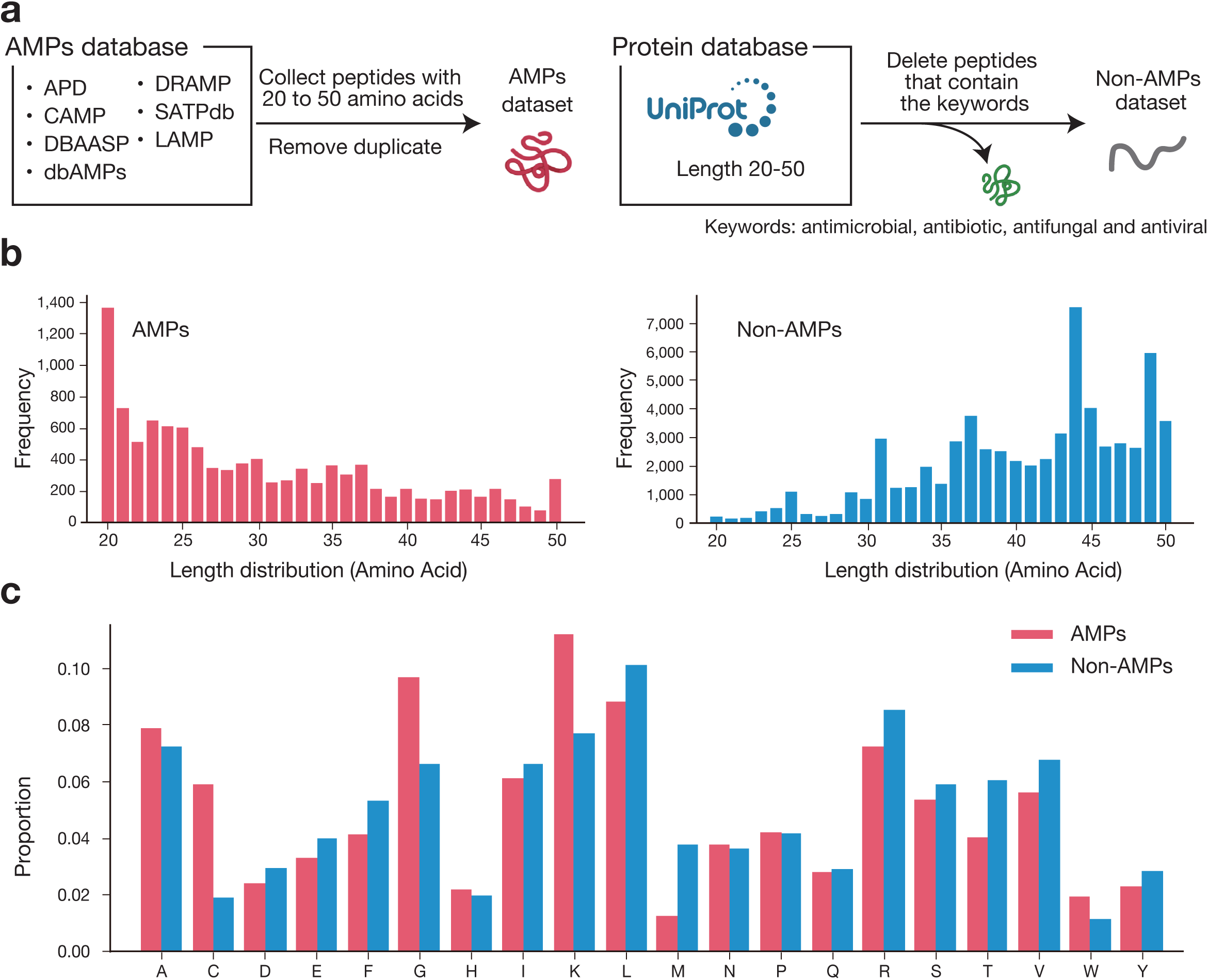
Details of the AMP-Atlas training dataset. **(a)** Method of data collection. AMPs sequence information was obtained from public databases by collecting AMPs with sequence lengths of 20–50 amino acids. Non-AMP sequence information was collected from UniProt, retrieving sequences of 20–50 amino acids that do not contain the keywords “antimicrobial,” “antibiotic,” “antifungal,” or “antiviral.” **(b)** Distribution of peptide sequence lengths in the training dataset (left: AMPs, right: non-AMPs). The horizontal axis represents the sequence length, and the vertical axis represents the frequency. **(c)** Amino acid composition in the training dataset (red: AMPs, blue: non-AMPs). The horizontal axis represents the type of amino acid, and the vertical axis represents the proportion of each amino acid.

## Reference

1. Murray, C. J. L. et al. Global burden of bacterial antimicrobial resistance in 2019: a systematic analysis. The Lancet 399, 629–655 (2022).

2. The Lancet. Antimicrobial resistance: an agenda for all. The Lancet 403, 2349 (2024).

3. Naghavi, M. et al. Global burden of bacterial antimicrobial resistance 1990– 2021: a systematic analysis with forecasts to 2050. The Lancet 404, 1199–1226 (2024).

4. Keiji Hirai, History and prospects of development of antimicrobial agents of Japanese origin. Japanese Journal of Chemotherapy 68, 499–509 (2020)

5. Piddock, L. J. The crisis of no new antibiotics—what is the way forward? Lancet Infect. Dis. 12, 249–253 (2012).

6. Geneva: & World Health Organization. 2023 Antibacterial agents in clinical and preclinical development: an overview and analysis. ISBN: 978-92-4-009400-0 (2024).

7. Reddy, K. V. R., Yedery, R. D. & Aranha, C. Antimicrobial peptides: premises and promises. Int. J. Antimicrob. Agents 24, 536–547 (2004).

8. Izadpanah, A. & Gallo, R. L. Antimicrobial peptides. J. Am. Acad. Dermatol. 52, 381–390 (2005).

9. Zhang, L. & Gallo, R. L. Antimicrobial peptides. Curr. Biol. 26, R14–R19 (2016).

10. Gan, B. H., Gaynord, J., Rowe, S. M., Deingruber, T. & Spring, D. R. The multifaceted nature of antimicrobial peptides: current synthetic chemistry approaches and future directions. Chem. Soc. Rev. 50, 7820–7880 (2021).

11. Ji, S. et al. Antimicrobial peptides: An alternative to traditional antibiotics. Eur. J. Med. Chem. 265, 116072 (2024).

12. Lazzaro, B. P., Zasloff, M. & Rolff, J. Antimicrobial peptides: Application informed by evolution. Science 368, eaau5480 (2020).

13. Chen, E. H.-L. et al. Visualizing the membrane disruption action of antimicrobial peptides by cryo-electron tomography. Nat. Commun. 14, 5464 (2023).

14. Huang, J. et al. Identification of potent antimicrobial peptides via a machine-learning pipeline that mines the entire space of peptide sequences. *Nat*. Biomed. Eng. 7, 797–810 (2023)

15. Fjell, C. D., Hiss, J. A., Hancock, R. E. W. & Schneider, G. Designing antimicrobial peptides: form follows function. Nat. Rev. Drug Discov. 11, 37–51 (2012).

16. Chen, S. et al. The discovery of antimicrobial peptides from the gut microbiome of cockroach Blattella Germanica using deep learning pipeline. http://biorxiv.org/lookup/doi/10.1101/2024.02.12.580024 (2024)

17. Torres, M. D. T., Melo, M. C. R., Crescenzi, O., Notomista, E. & De La Fuente-Nunez, C. Mining for encrypted peptide antibiotics in the human proteome. *Nat*. Biomed. Eng. 6, 67–75 (2021).

18. Wang, G., Li, X. & Wang, Z. APD3: the antimicrobial peptide database as a tool for research and education. Nucleic Acids Res. 44, D1087–D1093 (2016).

19. Sharma, K., Sharma, K. K., Sharma, A. & Jain, R. Peptide-based drug discovery: Current status and recent advances. Drug Discov. Today 28, 103464 (2023).

20. Asif, F., Zaman, S. U., Arnab, Md. K. H., Hasan, M. & Islam, Md. M. Antimicrobial Peptides as Therapeutics: Confronting Delivery Challenges to Optimize Efficacy. The Microbe 2, 100051 (2024).

21. Zhang, Q.-Y. et al. Antimicrobial peptides: mechanism of action, activity and clinical potential. Mil. Med. Res. 8, 48 (2021).

22. Pinacho-Castellanos, S. A., García-Jacas, C. R., Gilson, M. K. & Brizuela, C. A. Alignment-Free Antimicrobial Peptide Predictors: Improving Performance by a Thorough Analysis of the Largest Available Data Set. J. Chem. Inf. Model. 61, 3141–3157 (2021).

23. Veltri, D., Kamath, U. & Shehu, A. Deep learning improves antimicrobial peptide recognition. Bioinformatics 34, 2740–2747 (2018).

24. Li, C. et al. AMPlify: attentive deep learning model for discovery of novel antimicrobial peptides effective against WHO priority pathogens. BMC Genomics 23, 77 (2022).

25. Hideki Yamaguchi & Yutaka Saito, Protein language models, JSBi Bioinformatics Review, 4(1), 52–67 (2023).

26. Lin, Z. et al. Evolutionary-scale prediction of atomic-level protein structure with a language model. Science 379, 1123–1130 (2023).

27. Cordoves-Delgado, G. & García-Jacas, C. R. Predicting Antimicrobial Peptides Using ESMFold-Predicted Structures and ESM-2-Based Amino Acid Features with Graph Deep Learning. J. Chem. Inf. Model. 64, 4310–4321 (2024).

28. Wang, R. et al. Diff-AMP: tailored designed antimicrobial peptide framework with all-in-one generation, identification, prediction and optimization. Brief. Bioinform. 25, bbae078 (2024).

29. Li, K., et al. AMPCliff: quantitative definition and benchmarking of activity cliffs in antimicrobial peptides. Preprint at https://arxiv.org/abs/2404.09738 (2024)

30. Hu, E. J. et al. LoRA: Low-Rank Adaptation of Large Language Models. Preprint at http://arxiv.org/abs/2106.09685 (2021).

31. Jhong, J.-H. et al. dbAMP 2.0: updated resource for antimicrobial peptides with an enhanced scanning method for genomic and proteomic data. Nucleic Acids Res. 50, D460–D470 (2022).

32. Pirtskhalava, M. et al. DBAASP v3: database of antimicrobial/cytotoxic activity and structure of peptides as a resource for development of new therapeutics. Nucleic Acids Res. 49, D288–D297 (2021).

33. Shi, G. et al. DRAMP 3.0: an enhanced comprehensive data repository of antimicrobial peptides. Nucleic Acids Res. 50, D488–D496 (2022).

34. Singh, S. et al. SATPdb: a database of structurally annotated therapeutic peptides. Nucleic Acids Res. 44, D1119–D1126 (2016).

35. Zhao, X., Wu, H., Lu, H., Li, G. & Huang, Q. LAMP: A Database Linking Antimicrobial Peptides. PLoS ONE 8, e66557 (2013).

36. Gawde, U. et al. CAMPR4: a database of natural and synthetic antimicrobial peptides. Nucleic Acids Res. 51, D377–D383 (2023).

37. Xing, W., Zhang, J., Li, C., Huo, Y. & Dong, G. iAMP-Attenpred: a novel antimicrobial peptide predictor based on BERT feature extraction method and CNN-BiLSTM-Attention combination model. Brief. Bioinform. 25, bbad443 (2023).

38. Li, C. et al. Mining the UniProtKB/Swiss-Prot database for antimicrobial peptides. Preprint at 10.1101/2024.05.24.595811 (2024).

39. Das, P. et al. Accelerated antimicrobial discovery via deep generative models and molecular dynamics simulations. Nat. Biomed. Eng. 5, 613–623 (2021).

40. Zeng, Y. & Lee, K. The Expressive Power of Low-Rank Adaptation. Preprint at 10.48550/arXiv.2310.17513 (2024).

41. Aghajanyan, A., Zettlemoyer, L. & Gupta, S. Intrinsic Dimensionality Explains the Effectiveness of Language Model Fine-Tuning. Preprint at http://arxiv.org/abs/2012.13255 (2020).

42. Sohn, K. Improved Deep Metric Learning with Multi-class N-pair Loss Objective. NeurIPS (2016)

43. McInnes, L., Healy, J. & Melville, J. UMAP: Uniform Manifold Approximation and Projection for Dimension Reduction. Preprint at 10.48550/arXiv.1802.03426 (2020).

44. Cao, Q. et al. Designing antimicrobial peptides using deep learning and molecular dynamic simulations. Brief. Bioinform. 24, bbad058 (2023).

45. Wozniak, W. et al. Identification of human host factors required for beta-defensin-2 expression in intestinal epithelial cells upon a bacterial challenge. Sci. Rep. 14, 15442 (2024).

46. Wohlwend, J. et al. Boltz-1: Democratizing Biomolecular Interaction Modeling. Preprint at 10.1101/2024.11.19.624167 (2024).

47. Lehrer, R. I., Bevins, C. L. & Ganz, T. Defensins and Other Antimicrobial Peptides and Proteins. in Mucosal Immunology 95–110 (2005).

48. Petruschke, H. et al. Discovery of novel community-relevant small proteins in a simplified human intestinal microbiome. Microbiome 9, 55 (2021).

49. Hou, K. et al. Microbiota in health and diseases. Signal Transduct. Target. Ther. 7, 135 (2022).

50. Rakoff-Nahoum, S. & Medzhitov, R. Innate immune recognition of the indigenous microbial flora. Mucosal Immunol. 1, S10–S14 (2008).

51. Richardson, L. et al. MGnify: the microbiome sequence data analysis resource in 2023. Nucleic Acids Res. 51, D753–D759 (2023).

52. Hyatt, D. et al. Prodigal: prokaryotic gene recognition and translation initiation site identification. BMC Bioinformatics 11, 119 (2010).

53. Akiba, T., Sano, S., Yanase, T., Ohta, T. & Koyama, M. Optuna: A Next-generation Hyperparameter Optimization Framework. Preprint at http://arxiv.org/abs/1907.10902 (2019).

54. Loshchilov, I. & Hutter, F. Decoupled Weight Decay Regularization. Preprint at 10.48550/arXiv.1711.05101 (2019).

55. Leblanc, S. et al. OpenProt 2.0 builds a path to the functional characterization of alternative proteins. Nucleic Acids Res. 52, D522–D528 (2024).

56. Mirdita, M., Steinegger, M. & Söding, J. MMseqs2 desktop and local web server app for fast, interactive sequence searches. Bioinformatics 35, 2856–2858 (2019).

57. Rosowsky, A., Fu, H., Chan, D. C. M. & Queener, S. F. Synthesis of 2,4-Diamino-6-[2’-*O*-(ω-carboxyalkyl)oxydibenz[*b*,*f*]azepin-5-yl]methylpteridines as Potent and Selective Inhibitors of *Pneumocystis carinii, Toxoplasma gondii*, and *Mycobacterium avium* Dihydrofolate Reductase. J. Med. Chem. 47, 2475–2485 (2004).

58. Wells, B. A. & Chaffee, A. L. Ewald Summation for Molecular Simulations. J. Chem. Theory Comput. 11, 3684–3695 (2015).

59. Hoerschinger, V. J. et al. PEP-Patch: Electrostatics in Protein–Protein Recognition, Specificity, and Antibody Developability. J. Chem. Inf. Model. 63, 6964–6971 (2023).

60. Rego, N. B., Xi, E. & Patel, A. J. Identifying hydrophobic protein patches to inform protein interaction interfaces. Proc. Natl. Acad. Sci. 118, e2018234118 (2021).

